# Self-supervised contrastive learning for integrative single cell RNA-seq data analysis

**DOI:** 10.1101/2021.07.26.453730

**Authors:** Wenkai Han, Yuqi Cheng, Jiayang Chen, Huawen Zhong, Zhihang Hu, Siyuan Chen, Licheng Zong, Irwin King, Xin Gao, Yu Li

**Affiliations:** Computer, Electrical and Mathematical Sciences and Engineering Division (CEMSE), Computational Bioscience Research Center (CBRC), King Abdullah University of Science and Technology (KAUST), Thuwal, 23955, Saudi Arabia; Department of Computer Science and Engineering (CSE), The Chinese University of Hong Kong (CUHK), Hong Kong SAR, China; Weill Cornell Graduate School of Medical Sciences, Weill Cornell Medicine, New York, NY, 10065, USA; Biological and Environmental Sciences & Engineering Division (BESE), Red Sea Research Center (RSRC), King Abdullah University of Science and Technology (KAUST), Thuwal, 23955, Saudi Arabia; The CUHK Shenzhen Research Institute, Hi-Tech Park, Nanshan, Shenzhen, 518057, China

## Abstract

Single-cell RNA-sequencing (scRNA-seq) has become a powerful tool to reveal the complex biological diversity and heterogeneity among cell populations. However, the technical noise and bias of the technology still have negative impacts on the downstream analysis. Here, we present a self-supervised Contrastive LEArning framework for scRNA-seq (CLEAR) profile representation and the downstream analysis. CLEAR overcomes the heterogeneity of the experimental data with a specifically designed representation learning task and thus can handle batch effects and dropout events. In the task, the deep learning model learns to pull together the representations of similar cells while pushing apart distinct cells, without manual labeling. It achieves superior performance on a broad range of fundamental tasks, including clustering, visualization, dropout correction, batch effect removal, and pseudo-time inference. The proposed method successfully identifies and illustrates inflammatory-related mechanisms in a COVID-19 disease study with 43,695 single cells from peripheral blood mononuclear cells. Further experiments to process a million-scale single-cell dataset demonstrate the scalability of CLEAR. This scalable method generates effective scRNA-seq data representation while eliminating technical noise, and it will serve as a general computational framework for single-cell data analysis.

## Introduction

Single-cell RNA sequencing (scRNA-seq) has been a powerful tool for measuring the transcriptome-wide gene expression in individual cells and understanding the heterogeneity among cell populations^1, 2^. It has been facilitating researchers to investigate several critical biomedical topics, such as cancer^3^ and autoimmunity^4^. Despite its promises, the unique properties of the scRNA-seq data, such as extreme sparsity and high variability^5^, have posed a number of computational challenges to researchers^6, 7^. To analyze the data, among all the steps^7^, the key processing is to obtain a reliable low-dimension representation for each cell, which can preserve the biological signature of the cell for downstream analysis while eliminating technical noise^8, 9^.

The existing commonly used methods to perform the above processing are based on different backbone algorithms and assumptions. The earliest methods utilize the traditional dimension reduction algorithms, such as Principal Component Analysis (PCA), followed by *k*-means or hierarchical clustering to group cells^5, 10-15^. Although these methods are widely used, their assumption, that is, the complex single-cell transcriptomics can be accurately mapped onto a low-dimensional space by a generalized linear model, may not be necessarily justified^8^. Considering the complexity of the data, researchers have developed multiple kernel-based spectral clustering methods to learn more robust similarity matrices for cells^16, 17^. However, the time and space complexity of such methods impede the broad applications of the methods^5^. In contrast, the graph-based methods enjoy high speed and scalability^14, 15, 18, 19^. But such methods are hyper-parameter sensitive. The choice of *k* for the widely used *k*-nearest-neighbors graph affects the size and number of final clusters^7^. Because of the model capacity and scalability of deep learning methods, almost all the recently developed methods are based on antoencoder^5, 9, 20-25^ (AE) or variational autoencoder^8, 26, 27^ (VAE), which can also incorporate the biostatistical models^28, 29^ seamlessly. However, as AE and VAE methods are unsupervised learning methods, it is very difficult to control and decide what the deep learning models will learn, although some very recent studies try to impose constraints and our prior knowledge about the problem onto the low-dimensional space^5, 27^. Researchers have also tried to utilize manual labeling as supervision for training the models, accompanied by transfer learning^22^ or meta-learning^30^, but such methods encounter scalability issues and have strong assumptions on the homogeneity of different datasets, making them less popular than the above methods.

As discussed above, almost all the existing methods are based on unsupervised learning^7^, regardless of the specific algorithm. Without accessible supervision, for the deep learning-based methods, it is hard to guide the training process of the model and explain why a particular transformation is learned, although the model may work well. To promote the scRNA-seq data analysis, we indeed have some specific requirements for the model. For example, the functionally similar cells should be close in the transformed space, while distinct cells should be distant^7^; the model should overcome the batch effect and map the cells of the same type but from different experiments into the same region^8^. Unsupervised learning methods may have difficulty in incorporating these requirements explicitly. Here, we propose a novel method, CLEAR, for integrative single-cell RNA-seq data analysis, based on a new machine learning scenario, self-supervised learning, which can model all the above requirements explicitly. More specifically, we design our method based on self-supervised contrastive learning^31^, where we construct the training labels from the unlabeled data. For the gene expression profile of each cell, we distort the data slightly by adding noise to the raw data, which mimics the technical noise in the biological experiments. During training, we force the model to produce similar low-dimension representations for the raw data and the corresponding distorted profile (positive pairs). Meanwhile, we train the model to output distant representations for cells of different types (negative pairs). Intuitively, the deep learning model learns to pull together the representations of similar cells while pushing apart different cells, only utilizing labels constructed from the data without manual labeling.

Based on self-supervised contrastive learning, CLEAR achieves superior performance on a broad range of fundamental tasks for single-cell RNA-seq data analysis, including clustering, visualization, dropout correction, batch effect removal, and pseudo-time inference. As for clustering, CLEAR can outperform the popular tools and recently proposed tools on diverse datasets from different organisms. Applied on a dataset from a COVID-19 disease study with 43,695 single cells from peripheral blood mononuclear cells, CLEAR successfully identifies and illustrates inflammatory-related mechanisms. Further experiments to process a million-scale single-cell dataset demonstrate the scalability and potential of CLEAR to handle the emerging large-scale cell atlases. With the capability of generating effective scRNA-seq data representation while eliminating technical noise, the proposed method can serve as a general computational framework for single-cell data analysis.

## Results

### Overview of CLEAR

Unlike most existing methods, which are based on unsupervised learning to map the single-cell gene expression profile to the low-dimension space, we develop CLEAR base on self-supervised learning. That is, although we do not have the golden standard supervised information, such as the cell type, we train the deep learning model using the supervision constructed from the unlabeled data themselves. Notice that we can incorporate our prior knowledge about single-cell RNA-seq data, such as noise and dropout events, into the model training process implicitly and seamlessly when we build the label from the unlabeled data. More specifically, we design CLEAR based on self-supervised contrastive learning^31^. As shown in **Fig. 1**, eventually, we also want to train a deep learning encoder to map the gene expression profile into the low-dimension space. However, in addition to that, we further want the trained model to force functionally similar cells close in the transformed space while distinct cells being distant. Here, the model should also be robust to technical noise, such as dropout events. That is, the profiles from the same cell, no matter with or without dropout events, should be mapped into the same place in the low-dimension space. Although it is difficult to estimate the noise level of the real dataset, we can add simulated noise to the data and force the trained to be robust to them. Based on the above idea, we design CLEAR as shown in **Fig. 1**. Given the single-cell gene expression profile, we add different simulated noise, such as Gaussian noise and simulated dropout events, to it (data augmentation), resulting in distorted profiles (augmented data). The raw profile and the corresponding distorted profiles from the same cell are positive pairs, while the profiles from different cells are negative pairs. When training the model, we force the model to produce similar representations for the positive pairs while distinct for the negative pairs (contrastive learning). Intuitively, we pull together the representations of functionally close cells in the low-dimension space while pushing apart the embeddings of the dissimilar ones. CLEAR does not have any assumptions on the data distribution or the encoder architecture. It can eliminate technical noise and generate effective scRNA-seq data representation, which is suitable for a range of downstream applications, such as clustering, batch effect correction, and time-trajectory inference, as discussed below.

**Fig. 1.**
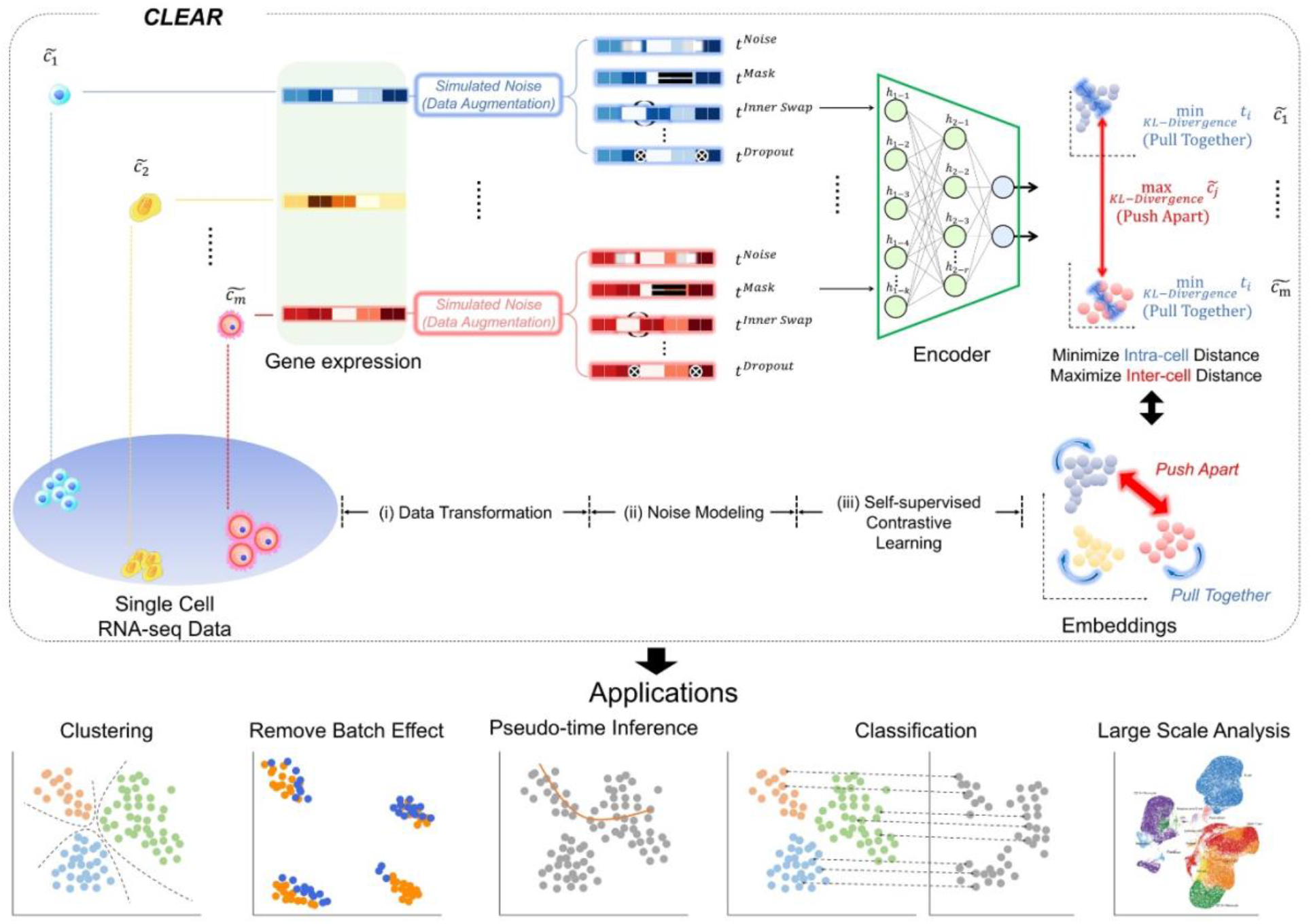
Overview of the proposed framework, CLEAR. The proposed method is based on self-supervised contrastive learning. For the gene expression profile of each cell, we distort the data slightly by adding noise to the raw data, which mimics the technical noise in the biological experiments. When training the deep encoder model, we force the model to produce similar low-dimension representations for the raw data and the corresponding distorted profile while distant representations for cells of different types. Intuitively, the deep learning model learns to pull together the representations of similar cells while pushing apart different cells. By considering noise during training, CLEAR can produce effective representations while eliminating technical noise for the scRNA-seq profiles. Such representations have a broad range of applications, including clustering and classification, dropout event and batch effect correction, pseudo-time inference. CLEAR is also scalable to million-scale datasets without any overhead.

### Overall clustering performance

To access how the representation from CLEAR helps to cluster, we evaluate the proposed method, combined with the *k* -means clustering algorithm, on ten published datasets with expert-annotated labels^32-38^. The label information is only available during testing. We compare our model with several state-of-the-art methods that are widely used for scRNA-seq data and belong to different categories, including PCA-based tools (Seurat^10^, SC3^11^, CIDR^12^, SINCERA^13^), graph-based methods (Seurat^18^, scGNN^24^), deep generative models (scVI^8^, scDHA^21^, scGNN^24^, ItClust^22^), and transfer learning approach (ItClust^22^). Evaluated on the same datasets with five-fold cross validation, CLEAR achieves substantially better performance in clustering adjusted Rand index (ARI) score than all the other methods on most datasets (**Fig. 2a, Supplementary Table 6)**. In particular, on average, CLEAR improves over the second-best method, scDHA, by 4.56% regarding the score. To evaluate the performance more comprehensively, we also use other metrics, such as normalized mutual information(NMI), where the compared methods show similar results. To better understand the representation produced by each method, we use uniform manifold approximation (UMAP) to project the internal representations into a two-dimensional space and visualize them. (**Fig. 2b, Supp Fig 1-9**) As shown in the figure, CLEAR learns to embed similar cells within the same clusters while separate dissimilar cells well among different clusters. Compared to the other methods, it produces more similar clustering results as the ground truth cell annotation. Furthermore, as illustrated in **Fig. 2c**, the river plot of the Hravtin dataset, comparing the CLEAR clustering and expert annotation, suggests that they are nearly perfectly matched. On the other hand, scDHA tends to under-cluster the dataset, *e*.*g*., the interneurons are mixed up with the Exicitory cell, while Seurat is likely to over-cluster the cells, *e*.*g*., oligodendrocytes and Excitatory cells are split into many subclusters. Although CLEAR does not access any human supervision on marker genes, it can recover the ground truth directly for this dataset, suggesting that the proposed framework can implicitly capture the data’s biological features. Furthermore, to demonstrate the effect of the proposed self-supervised contrastive learning settings, we perform an ablation study on the data augmentation operations, removing each operation one by one and recording the performance change. As shown in **Supplementary Table 8,10**, removing either augmentation step will lead to the decreased ARI performance of CLEAR, which strongly indicates that the introduced noise and instance discrimination task help the model to capture the real cell-cell relationships. In addition, we further check the influence of using highly variable genes, discovering that those low variable genes may reduce the signal ratio and harm the model performance, which is consistent with the previous study^24, 39^. The design and comprehensive results of the ablation studies, together with the hyper-parameter selection, are detailed in the **Supplementary Table 8-12**.

**Fig. 2.**
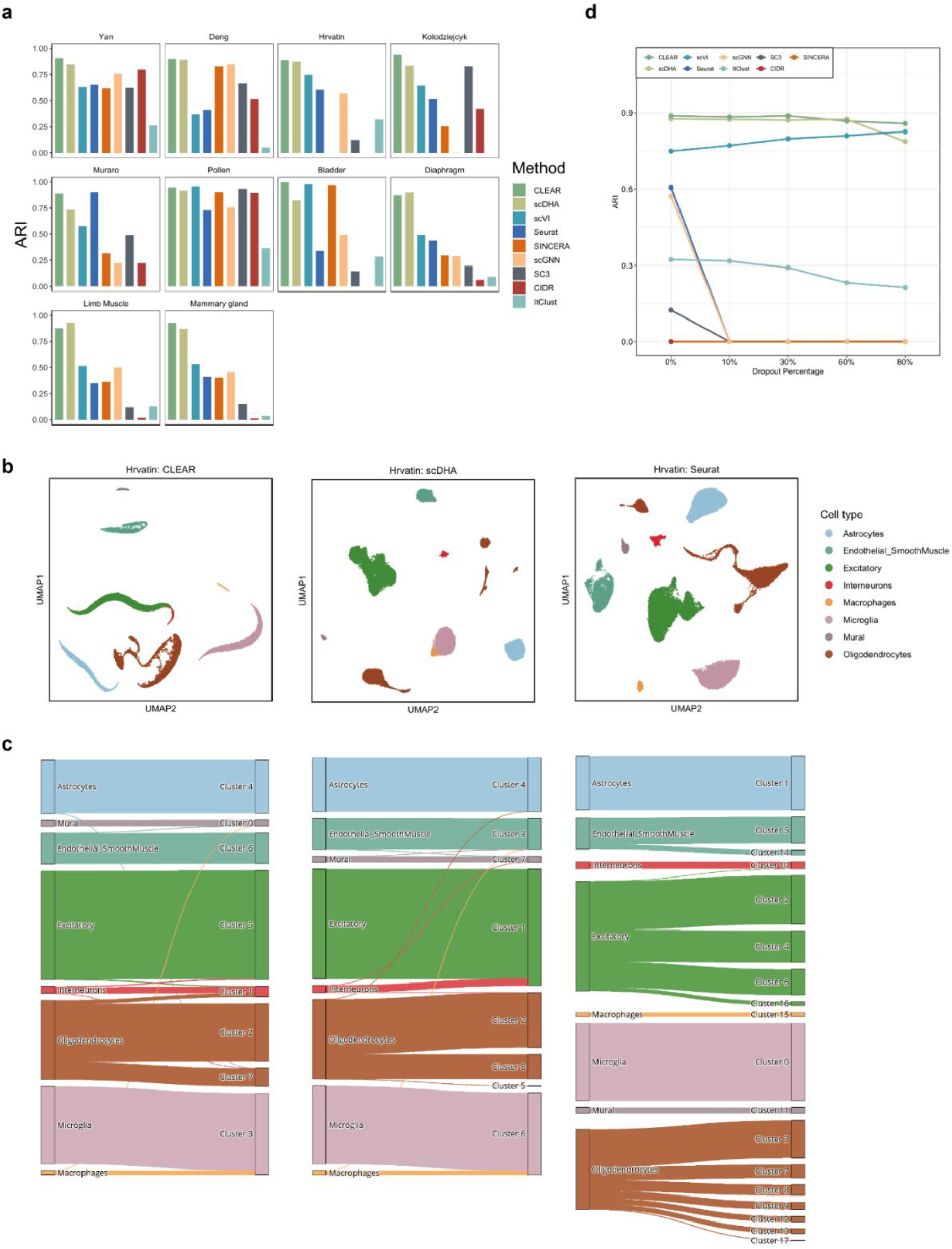
The representation from CLEAR benefits clustering and dropout event correction. **a** Clustering performance comparison of different methods on diverse datasets. On average, CLEAR improves over the second-best method, scDHA, by 4.56%, regarding ARI. **b** UMAP visualization of representations produced by CLEAR, scDHA, and Seruat on the Hrvatin dataset. **c** River plots of the Hrvatin dataset. CLEAR clustering matches almost perfectly with the expert annotation, without over-clustering or under-clustering. **d** Clustering performance change of different methods against different artificial dropout percentages in terms of ARI.

### CLEAR corrects dropout events and batch effects effectively

Dropout events and batch effects are notorious in scRNA-seq data analysis, which should be handled properly. We next evaluate the robustness of CLEAR when encountering dropout events. Although it is impossible to recover the actual gene expression levels and determine how dropouts impact the data, we simulate the dropout effects by randomly masking non-zero entries into zero with a hypergeometric distribution. Given the additional artificial dropouts, clustering becomes much more difficult. We test the eight competing approaches together with CLEAR on the Hravtin dataset, containing 48,266 single cells with 25,187 genes and thus 1.2 billion read counts. Among these reads, 94.2% of them are zeros. We set 10%, 30%, 60%, 80% dropout rates for the non-zero entries, respectively, resulting in a masked dataset with up to 98.8% zeros (**Supp Fig. 16**). CLEAR achieves the best performance in handling dropout events in terms of clustering, even when 80% of the non-zero entries are masked, suggesting that it is robust and has the potential to extract important features in some extreme cases. Although the performance of scDHA is similar to that of CLEAR when no dropouts are introduced, it becomes worse when the dropout rate is 80%. (**Fig. 2d**).

Although several methods have been proposed to correct batch effects, which are undesirable variability in the scRNA-seq datasets from technical and biological noise, most of them work as separate modules, focusing on one variable, and thus cannot generalize to the large complex atlas projects. CLEAR, however, has the potential to model multiple batch effects in an end-to-end fashion. Here, we assess CLEAR on correcting batch effects. Specifically, we first evaluate CLEAR on a dataset^40^, consisting of batches with shared cell types and biologically similar but unshared cell types. The goal of the batch effect removal algorithms is to integrate common cell types while maintaining separation between highly similar cells in different batches (**Methods**). As shown in **Fig. 3a**, CLEAR can separate difference cell types while mix up DoubleNeg and pDC cells from different batches. The biological similarity between CD141 cells and CD1C cells is also represented on the figure: the distance between CD141 cell cluster and CD1C cell cluster is closer than the other two clusters. On the other hand, scVI and SIMLR bring DoubleNeg and pDC cells closer but do not mix the batches well. Seurat can mitigate the batch effects in DoubleNeg and pDC cells but split CD141 cells into 2 clusters. ItCLUST mixed up all cells, regardless of batch and cell type, suggesting that it could not handle the dataset.

**Fig. 3.**
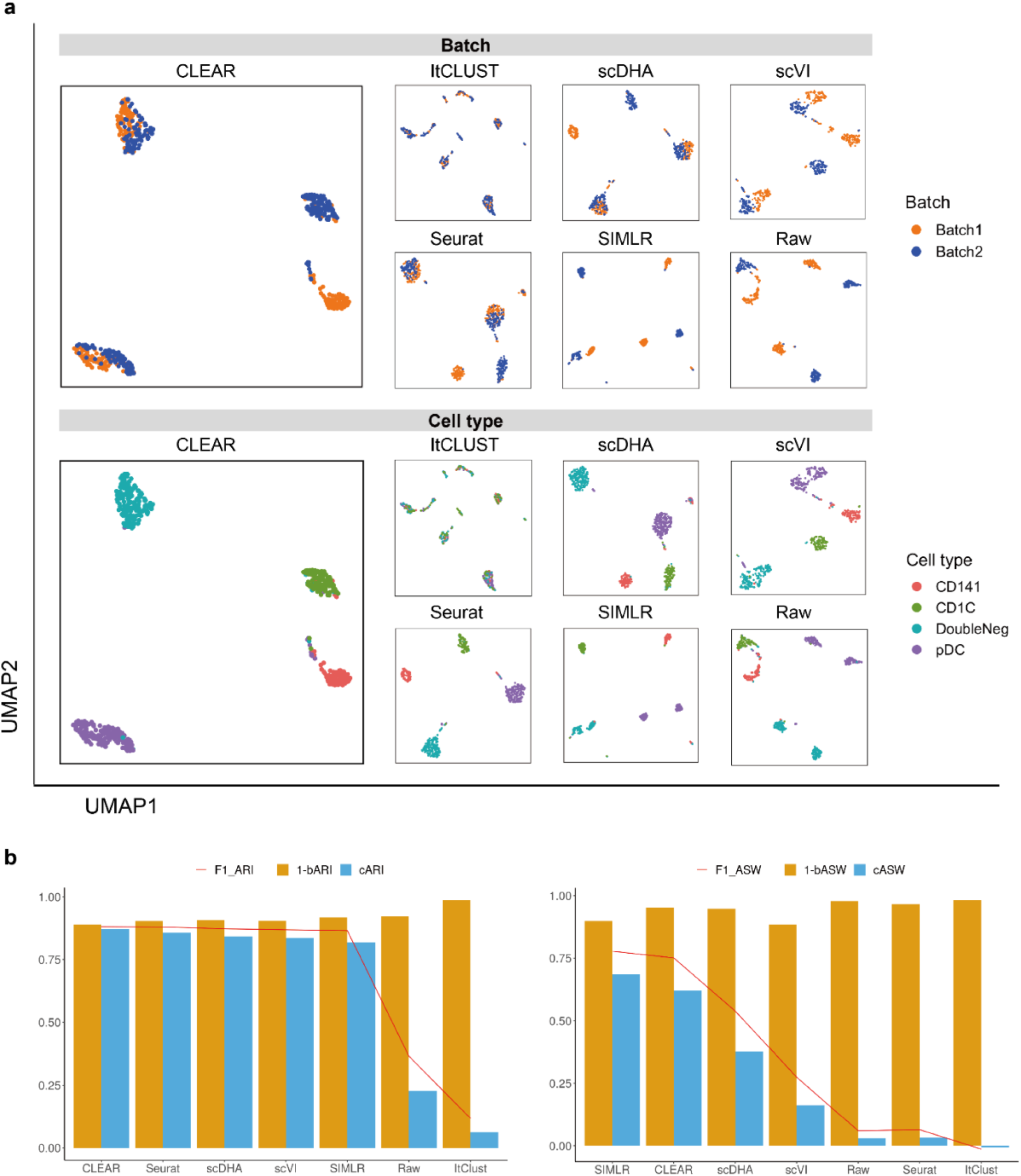
CLEAR corrects batch effects effectively. **a Upper panel:** UMAP visualization showing different methods’ performance on integrating DoubleNeg and pDC cells from two batches. **Bottom panel:** UMAP visualization showing different methods’ performance on separating four cell types. Notice that CLEAR’s representations also preserve the biological similarity between CD141 cells and CD1C cells. **b** The quantitative performance of different methods on batch effect removal, measured by adjusted rand index(ARI) and average silhouette width (ASW).

We further quantify the performance of different methods regarding batch effect removal with two metrics, average silhouette width (ASW) and adjusted rand index(ARI), on six datasets (**Datasets**). We further calculate each metric in three aspects: cell type (cARI, cASW), batch mixing (1 − *bARI*, 1 − *bASW*), and the Harmonic mean of the two (f1_ARI, f1_ASW). As shown in **Fig. 3b**, CLEAR achieves the best balance between cell type identification and batch mixing. Futhermore, CLEAR outperforms all the other baselines under various complex batch effects settings, even though it was not designed to do so (**Supp Fig. 10-15**). In particular, on the Tabula Muris Senis cell atlas, which covers the life span of a mouse and contains many batches, including cells from several mouses with different identities, ages, genders, and from different chips, CLEAR mixes all the cells of the same type from different batches while separating distinct cell types well.

### Pseudo-time inference

Another thriving topic in single-cell RNA-seq data analysis is pseudo-time inference, also known as trajectory inference. It aims to infer the ordering of cells along a one-dimensional manifold (pseudo-time) from the gene expression profiles. Usually, the inferring algorithms will benefit much from better data representations. Here, we evaluate whether the representation produced by CLEAR can facilitate the downstream pseudo-time inference. We use the CLEAR embeddeings and the PAGA^41^ algorithm to generate the pseudo-time. We compare it with two other popular methods, SCANPY^14^ and Monocle3^42^, using two mouse embryo development datasets: Yan^33^ and Deng^32^. We show the cells ordered by pseudo-time in **Fig. 4**. Ideally, the points should fall on the diagonal, indicating the relative relationship among the cells. The time inferred with CLEAR is strongly correlated with the true development stages, where Monocle3 mixes the cells from different development stages. We also use the R-squared value to quantify the performance. CLEAR achieves the highest value (R^2^ = 0.957), compared with SCANPY (R^2^ = −0.014) and Monocle3 (R^2^ = 0.884). We further illustrate the cell embeddings in the 2D space with UMAP, as shown in **Fig. 4**. The smooth lines indicate the time-trajectory from different methods. The trajectories inferred by CLEAR follow the development stages precisely. It starts at the zygote, goes through two cells, four cells, eight cells, and 16 cells, and finally stops at the blast cells. However, for Monocle3 and SCANPY, there is no clear trajectory among the cells. The cells in the early stages tend to mix, while cells in the late stages form another big group. The above experiments suggest that cell embeddings from CLEAR can facilitate the downstream algorithms in producing better biologically meaningful trajectories.

**Fig. 4.**
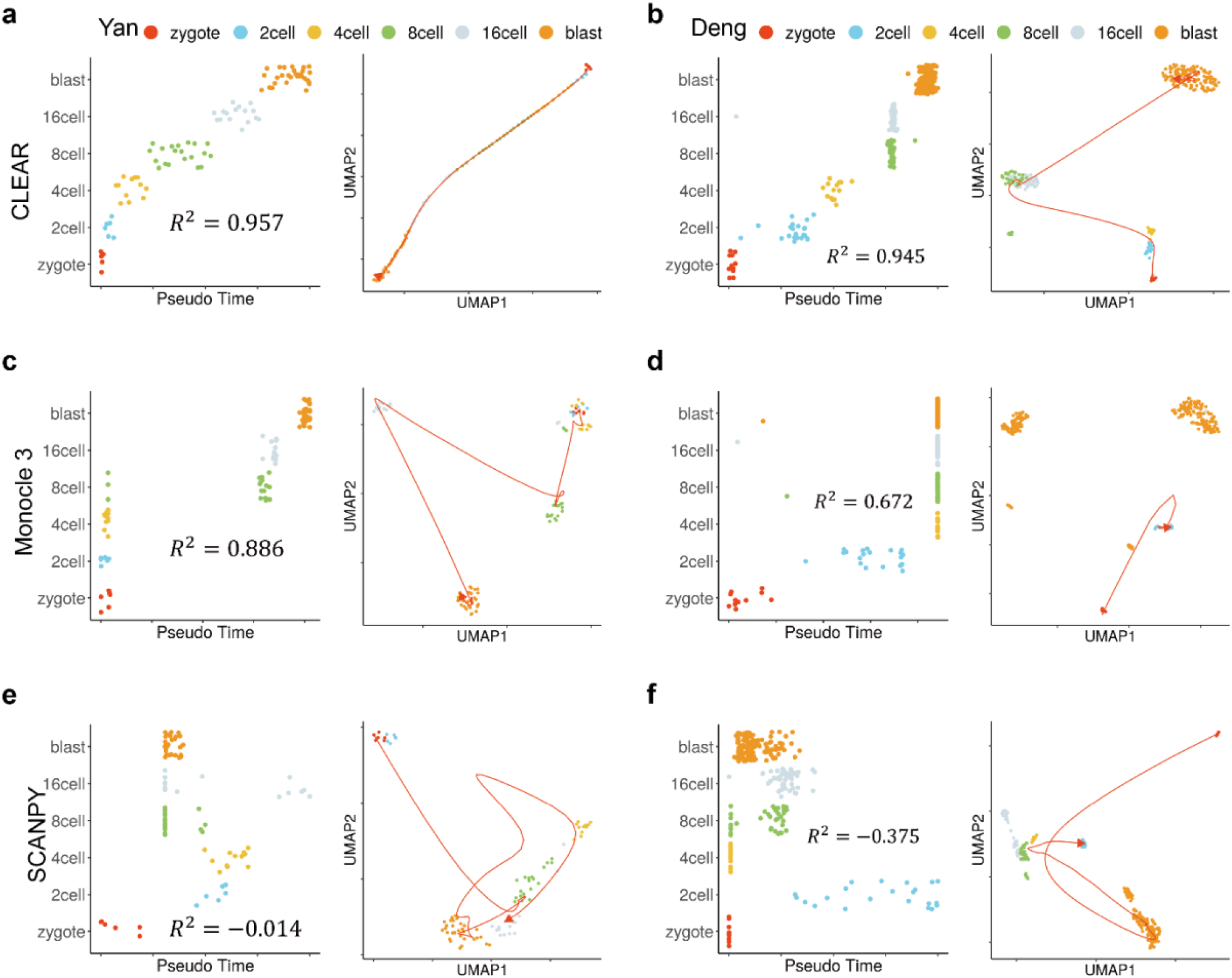
CLEAR is helpful for pseudo-time inference. **a** CLEAR’s performance on the pseudo-time inference for the Yan dataset. **Left figure:** Cells from the Yan dataset ordered by pseudo-time inferred from CLEAR. Ideally, the points should fall on the diagonal. **Right figure:** UMAP visualization of time trajectory inferred from CLEAR. **b** CLEAR’s performance on the pseudo-time inference for the Deng dataset. **c,d** Monocle3’s performance on the pseudo-time inference for the Yan and Deng dataset. **e,f** SCANPY’s performance on the pseudo-time inference for the Yan and Deng dataset.

### CLEAR illustrates peripheral immune cells atlas and inflammatory-related mechanisms in COVID-19

To demonstrate the application potential of CLEAR on real-world biology research, we apply it to analyze a newly published COVID-19 dataset^43^ (GEO accession number GSE150728), containing 44721 cells (43695 cells after quality control) collected from six healthy controls and seven COVID-19 samples. Four of the seven COVID samples are collected from patients with acute respiratory distress syndrome (ARDS) in clinical (**Fig. 5a, Supplementary Table 1**). We perform dimensionality reduction by CLEAR and graph-based clustering, identifying 32 clusters and visualizing them via uniform manifold approximation and projection (UMAP). We calculate the differential expressed genes (DEGs) of each cluster to annotate cell types manually. The cell types of monocytes (CD14+ and CD16+), T cells (CD4+ and CD8+), natural killer (NK) cells, B cells, plasmablasts, conventional dendritic cells (DCs), plasmacytoid dendritic cells (pDC), stem cell (SC) and eosinophil, neutrophil, platelets, and red blood cells (RBCs) are identified (**Fig. 5b,d, Supplementary Table 2**).

**Fig. 5.**
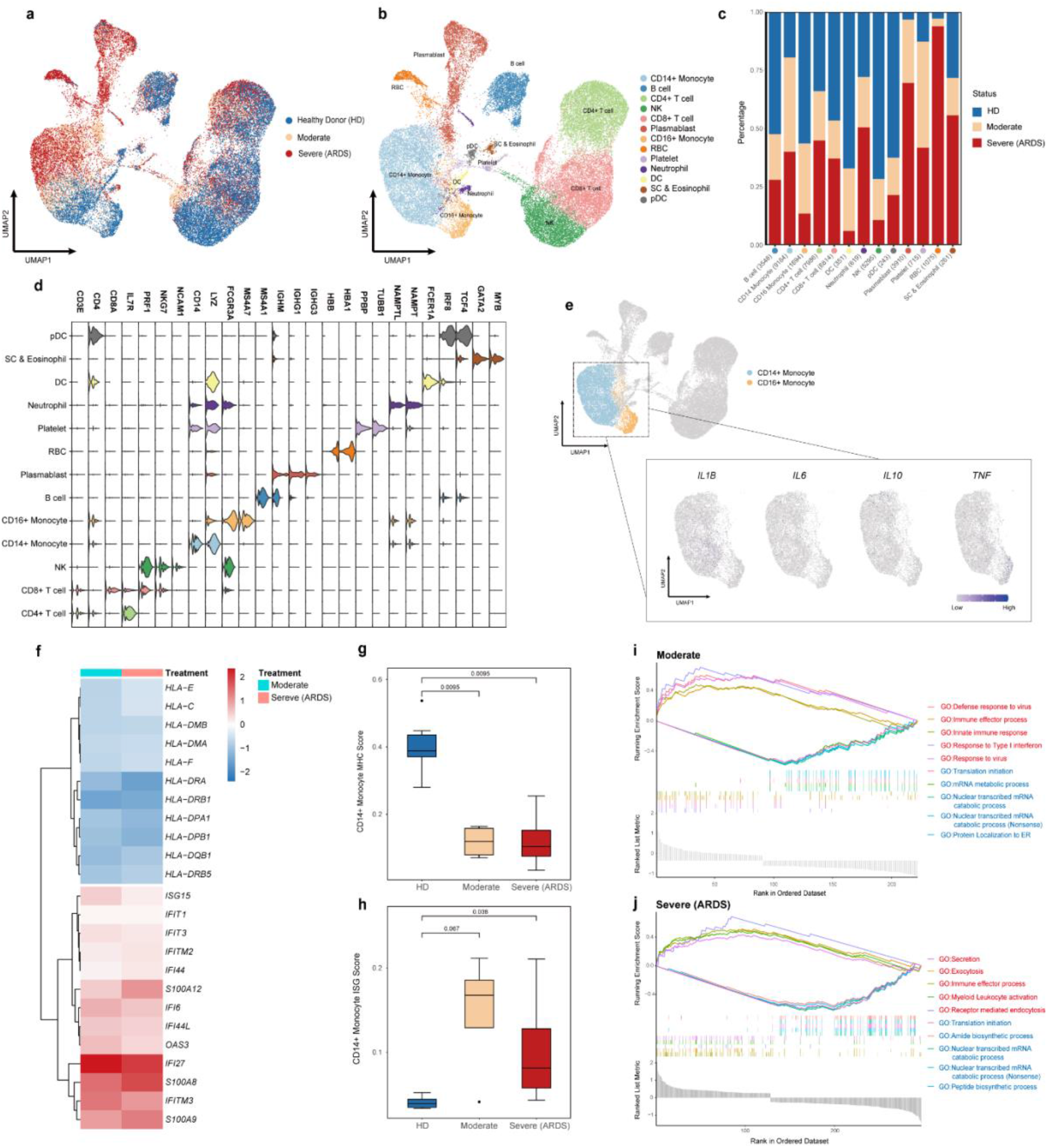
Peripheral immune cells atlas and inflammatory-related mechanisms in COVID-19 revealed by CLEAR. **a,b** UMAP visualization of the COVID-19 cell atlas **(a)** colored by COVID status and **(b)** colored by 13 cell type clusters (n=43695 cells). **c** Bar plot showing the relative percentage of different cell types comparing three COVID-19 statuses (HDs, Moderate status, and Severe status). **d** Stacked violin plot overview of the top-important marker genes expression for each cell type. **e** UMAP visualization of the key pro-inflammatory cytokines expression in both CD14+ and CD16+ monocytes. **f** Heatmap of IFN-stimulated genes and MHC-related genes in CD14+ monocyte. **g,h** Boxplots showing the mean **(g)** MHC-related score and **(h)** ISG score in CD14+ monocyte colored by different COVID statuses (HDs--blue, Moderate--Oranger and Severe (ARDS)--Red). **i,j** Gene set enrichment analysis (GSEA) of differential expressed gene (LogFC > 0.25) sets between **(i)** moderate CD14+ monocyte and healthy donor CD14+ monocyte and **(j)** severe CD14+ monocyte and healthy donor CD14+ monocyte. Red represents upregulated GO biological pathway, and blue represents downregulated GO biological pathway.

To assess the general atlas of immune responses and perturbation during different COVID-19 statuses, we quantify the proportions of immune cell subsets in health donors (HDs), moderate (without ARDS), or severe COVID-19 (with ARDS) individuals (**Fig. 5c**). Consistent with previous reports^43-45^, several immune cell subsets vary between healthy donors and COVID-19 samples, and we observe a significant depletion of NK cells, DC, pDC, and CD16+ monocytes. We also note an elevated frequency of plasmablasts, especially in patients with ARDS, which indicates that, together with the published clinical observations^46^, acute COVID-19 response may be associated with a severe humoral immune response.

Several previous studies have shown that severe COVID-19 has been associated with dysregulated immune responses, which may be induced by the abnormal activation or suppression of inflammatory reaction^47-50^. To reveal inflammatory-related mechanisms in COVID-19, we perform transcription level analysis on monocytes in more granularity. We first examine the expression of ‘COVID cytokine storm’ marker genes which encode pro-inflammatory cytokines reported before produced by monocytes, including *IL1B, IL2, IL6, IL10, TNF*^51, 52^. Interestingly, we do not find significant expression of these pro-inflammatory genes in monocytes (**Fig. 5e**), consistent with recent research with deeper profiling of immune cells^43, 49^, suggesting that COVID-19 may also present an immune suppression status. To further analyze transcription changes driving monocyte response remodeling in COVID-19, we conduct differential expression (DE) analysis and cellular pathway analysis by comparing COVID samples to HDs. Given that the dysregulation of CD14+ monocyte plays a more dominant role in COVID-19 progress^53^, we especially investigate the transcription profile changes in CD14+ monocytes. An increased IFN-stimulated gene (ISG) set and decreased major histocompatibility complex (MHC) molecules in CD14+ monocyte compared to HDs are observed (**Fig. 5f)**. Scoring the samples with published MHC-related genes and ISGs respectively also reveal that downregulation of MHC gene expression and upregulation of ISGs are significant in CD14+ monocytes across all the COVID patients (**Fig. 5g, h**). The dominant effect of the IFN response is consistent with the acute viral infection. But the suppression of MHC molecules may hinder the ability to activate lymphocytes and raise an effective anti-viral response. We then apply Gene Ontology (GO) analysis, combined with GSEA, to study the biological pathway changes in CD14+ monocytes with different COVID statuses. Significant ISG upregulation in CD14+ monocyte in moderate samples is also reflected in the pathway analysis, such as Type I interferon response (**Fig. 5i**), which may indicate a more active interferon level in moderate COVID patients and have the potential to become a clinical blood test marker to monitor COVID progress. Interestingly, we also find a secretion pathway and myeloid leukocyte activation upregulation in severe samples (**Fig. 5j**). This may suggest a dysregulated CD14+ monocytes activation in patients with ARDS.

### CLEAR handles million-scale scRNA-seq datasets

With the unprecedented increase in sequencing scale of the recent scRNA-seq experiment platform, the ability to process million-scale single-cell sequencing datasets is increasingly essential. However, many published tools require complicated parameter setting tunning and cause burdens on the users with the split-merge process^54^. This has become a big challenge. The proposed method, CLEAR, is a robust and scalable framework, which can resolve the problem naturally. It can perform million-level dataset dimension reduction in parallel while getting rid of the tedious parameter tunning process. To test the scalability of CLEAR, we apply it on a newly published million-level COVID PBMC scRNA-seq dataset (GEO accession number GSE158055), which contains around 1.5 million cells from COVID samples. We use CLEAR with the default parameters to conduct dimension reduction, visualizing the produced representations of the dataset with UMAP. CLEAR identifies 40 clusters, which are then annotated manually according to each cluster’s top 100 differential expressed genes (**Fig. 6a**). Among them, we find 14 subtypes and then plot selected marker genes for each cell type. Satisfyingly, a significant expression track of these marker genes is obtained under the higher level (*e*.*g*., CD4+ T cells and CD8+ T cells are combined as T cells) of these subtypes (**Fig. 6b**). Performing sensitive feature extraction while eliminating technical noise on the million-scale dataset, CLEAR is an easy-to-use and well-performed large-scale scRNA-seq data analysis tool, which has the potential to assist the construction and refinement of cell atlases.

**Fig. 6.**
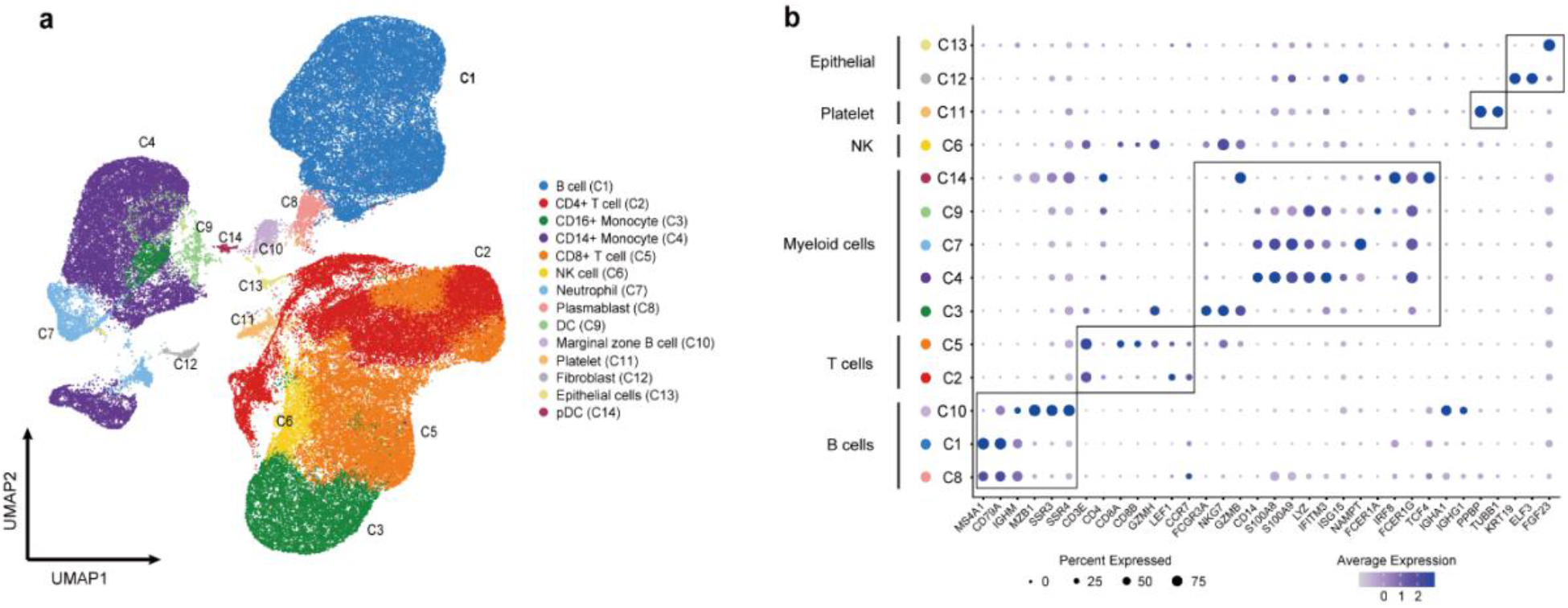
COVID-19 PBMC cell atlas based on million-scale scRNA-seq dataset. **a** UMAP embedding of peripheral blood mononuclear cells (PBMCs) from all samples (n=1.46 million cells) colored by manually-added cell types. **b** Dot plot showing percent expression and average expression of the selected marker genes for each cell type.

## Discussion and Conclusion

scRNA-seq has become a powerful and essential tool in biological research. With the accumulated data and the emerging cell atlases, the demand for practical computational tools to process and analyze such data has never been fully satisfied. Based on the current situation, the newly developed tools to process the single-cell data should, first of all, learn effective representations for the profile while eliminating the technical noise within the data. Secondly, they should have sufficient scalability to handle the million-scale unlabeled datasets in the field. Here, we introduce such a framework, CLEAR, based on self-supervised contrastive learning. By introducing noise during training and forcing the model to pull together the representation of functionally similar cells while pushing apart dissimilar cells with a carefully designed task, we managed to train the model to produce effective representations for the single-cell profile. CLEAR achieves superior performance on a broad range of fundamental tasks, including clustering, visualization, dropout correction, batch effect removal, and pseudo-time inference. Furthermore, it scalable enough to handle a million-scale dataset, which suggests its potential to handle the emerging cell atlases.

In the future, CLEAR can be further developed from both the biological aspect and the machine learning aspect. Regarding the biological application, obviously, CLEAR is a very flexible framework to perform data integration, no matter the single-omics, multi-dataset integration (cell atlases construction), or multi-omics integration (e.g., the integration of scRNA-seq and scATAC-seq data). In terms of the machine learning technical details, more advanced methods to handle data imbalance and incorporate prior knowledge, such as partially labeled data, should be developed. We believe that our framework, CLEAR, will become an alternative approach for single-cell data analysis.

## Methods

### Datasets

Below we describe how we obtained and preprocessed each dataset. Notice that, to show the generalization property of CLEAR, we used various datasets with different sequencing protocols (Smart-Seq, Smart-Seq2, DropSeq, CEL-Seq2, *etc*.), from different tissues, and with diverse data sizes (from 90 cells to 1.46 million cells). Unless otherwise noted, we obtained the cell type annotation information of each dataset from the original data paper. For the Hrvatin, Kolodziejczyk, Muraro, Pollen, and Tabula Muris Senis datasets, to improve the data quality, we filtered out low-quality cells with fewer than 200 genes and genes expressed in less than three cells.

Yan dataset. The Yan dataset refers to the human preimplantation embryos and embryonic stem cells. In this dataset, 90 cells were sequenced with the Tang protocol. We downloaded the dataset from Hemberg Group’s website. We used Scanpy to log-transform the dataset, and then each cell was normalized to 10,000 read counts. After that, highly variable genes were selected. Finally, we scaled the dataset to unit variance and zero mean.

Deng dataset. The Deng dataset refers to the mouse preimplantation embryos and embryonic stem cells of mixed background. 268 cells were sequenced via two protocols, Smart-Seq and Smart-Seq2. We downloaded the dataset from Hemberg Group’s website and performed a log-normalization transformation on the RPKM expression values with SCANPY.

Hrvatin dataset. The Hrvatin dataset refers to the mouse primary visual cortex cells under different simulation conditions, which was downloaded from GEO database (accession number: GSE102827). It contains 48,266 cells from 6-8 week-old mice, which were sequenced by DropSeq. After low-quality data filtering, we performed the log transformation, per-cell count normalization, and highly variable gene selection steps as mentioned above.

Kolodziejczyk dataset. This dataset, which was downloaded from the Hemberg Group’s website, contains 704 embryonic stem cells. They were sequenced with three batches under SMARTer protocol. We removed the low-quality data and the spiked-in cells. The top 2,000 most variable genes were selected for downstream analysis after we normalized each cell to 10,000 read counts.

Muraro dataset. This dataset, sequenced with the CEL-Seq2 protocol, contains 2126 cells from the human pancreas. We downloaded the data from the GEO database (accession number GSE85241), and removed the low-quality data and cells containing a higher number of mitochondrial genes and spike-in RNAs. The highly variable genes were then selected after log transformation.

Pollen dataset. The dataset, sequenced with the SMARTer protocol, contains 301 cells in the developing cerebral cortex from 11 populations. We downloaded it from the Hemberg Group’s website, removing the low-quality data and cells having a higher number of mitochondrial genes and spike-in RNAs. The highly variable genes were then selected after log transformation.

Tabula Muris Senis dataset. The entire dataset, sequenced with 10X, contains more than 100,000 cells. It was generated across the lifespan of mice, including 23 tissues and organs, but here we only focus on four tissues: bladder, mammary gland, limb muscle, and diaphragm. These datasets allow us to examine the batch effect and cell clustering. After downloading the raw data from https://figshare.com/projects/Tabula_Muris_Senis/64982, we removed the low-quality data and cells containing a higher number of mitochondrial genes and spike-in RNAs. Each cell was normalized to 10,000 read counts. The highly variable genes were then selected after log transformation. Finally, we scaled the dataset to unit variance and zero mean.

Human dendritic cells dataset. It consists of human blood dendritic cell (DC) data from Villani *et al* ^*55*^. We downloaded the data from https://github.com/JinmiaoChenLab/Batch-effect-removal-benchmarking/tree/master/Data. The dataset is composed of two batches. Each batch contains three cell types. Both batches share two cell types (pDC and double negative), while remaining one unshared biologically similar cell type (CD141 and CD1C, respectively).

COVID PBMC dataset. This dataset (GEO accession number: GSE150728), generated by Wilk *et al* ^*43*^, contains 44,271 cells sequenced with the Seq-Well platform. We have eight peripheral blood samples from seven SARS-COV-2 patients and six healthy controls. We removed the cells with a higher number of mitochondrial genes and spike-in RNAs. Each cell was normalized to 10,000 read counts. The highly variable genes were then selected after log transformation. Finally, we scaled the dataset to unit variance and zero mean. The cell type information was annotated using marker genes by experts.

COVID large-scale dataset. This dataset (GEO accession number: GSE158055), generated by Ren *et al* ^*56*^, contains more than 1.46 million cells generated through 10X Genomics. 171 COVID-19 patients and 25 healthy individuals were enrolled with PBMC, BALF, PFMC, and sputum samples. We removed the cells containing a higher number of mitochondrial genes and spike-in RNAs. Each cell was normalized to 10,000 read counts. The highly variable genes were then selected after log transformation. Finally, we scaled the dataset to unit variance and zero mean. The cell type information was annotated using marker genes by experts.

### The CLEAR framework

The key idea of CLEAR is to learn effective cell representations, considering noise in the data, and to pull together the representation of functionally similar cells, while pushing apart dissimilar cells. We achieve the goal with self-supervised contrastive learning. Given the single-cell gene expression profile, we add different simulated noise, such as Gaussian noise and simulated dropout events, to it (data augmentation), resulting in distorted profiles (augmented data). The raw profile and the corresponding distorted profiles from the same cell are positive pairs, while the profiles from different cells are negative pairs. When training the model, we force the model to produce similar representations for the positive pairs while distinct for the negative pairs (contrastive learning). More specifically, by discriminating the positive pairs from a large number of negatives, CLEAR learns a locally smooth nonlinear mapping function *f*_*θ*_ that pulls together multiple distortions of a cell in the embedding space and pushes away the other samples. The locally smooth function is also helpful for the global embeddings. In the transformed space, cells with similar expression patterns form clusters, which are likely to be cells of the same cell types. The function *f*_*θ*_ is parameterized by a deep neural network, whose parameters can be optimized in an end-to-end manner. The detailed workflows are as below.

1. Data augmentation. We first perform data augmentation to generate training pairs. Each cell will have two augmented versions, and thus a minibatch of *N* cells is augmented to 2*N* cells. This step will be discussed in detail in **Data augmentation**.
2. Constructing negative labels with data from multiple minibatches. For data in one minibatch, we can consider the two data points generated from the same gene expression profile as a positive pair while the other combinations as negatives. However, if we only consider the negatives within a minibatch, the learned mapping function is less likely to be effective for global clustering. To make the locally smooth function *f*_*θ*_ have a global effect, we should consider negatives from other minibatches. We achieve that by maintaining a queue with data from multiple minibatches. When the current minibatch is enqueued, the oldest minibatch will be dequeued. Within the queue, a specific distorted profile only has one positive pair match, while all the other profiles are negatives for it. Notice that the conceptual difference between minibatch and the queue arises from the hardware limitation. If the GPU memory is large enough and we can feed all the data in one minibatch, we can discard the queue maintenance.
3. Loss function. Let 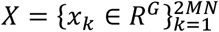 be the queue consisting of a number of gene expression profiles, where *G* denotes for the number of genes; *N* stands for the batch size; *M* stands for the number of batches stored in the queue. In one batch, *N* samples are augmented into 2*N* samples. Consequently, the queue consisting of *M* minibatches contains 2*MN* augmented samples. *x*_*k*_ denotes for the *k* -th (distorted) gene expression profile in the queue. For a pair of positive samples *x*_*i*_ and *x*_*j*_ (derived from one original sample), the other 2*MN* − 2 samples are treated as negatives. To distinguish the positive pair from the negatives, we use the following pairwise contrastive InfoNCE loss:

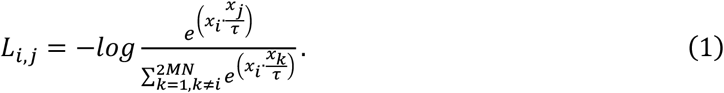

Note that *L*_*i,j*_ is asymmetrical. Suppose we put all the pairs in an order, such that 2*i* − 1 and 2*i* denote for the paired augmentations, then the summed-up loss is:

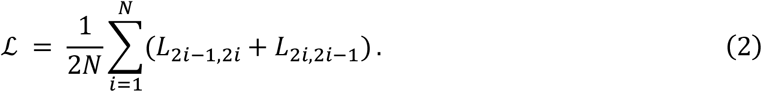
4. Momentum update. As suggested by He *et al*. ^57^, a rapidly changing encoder network will reduce the representations’ consistency, resulting in poor performance. To deal with the problem, we utilize two encoders, a slow-evolving key encoder *f*_*k*_ and a fast-evolving query encoder *f*_*q*_. Denoting the parameters of *f*_*q*_ as *θ*_*q*_ and those of *f*_*k*_ as *θ*_*k*_, we update the query encoder by the normal back-propagation. However, for the key encoder, we update it with momentum:

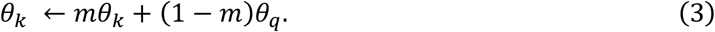

Here *m* ∈ [0,1) is a momentum coefficient. The momentum update makes the encoder network evolve smoothly.
5. Inference. After we train the model, the query encoder network *f*_*q*_ is the final productive network, which outputs the representation of a single cell gene expression profile. After obtaining the representations of all the cells in a dataset, we cluster the cells with the common clustering algorithms (*e*.*g*., k-means algorithm, Louvain algorithm, and Leiden algorithm). Finally, cell types are assigned to the discovered clusters based on the differential expression genes in the cluster.

### Data augmentation

Data augmentation is critical to the success of self-supervised contrastive learning. We use the following ways of data augmentation, considering noise during real experiments. Note that the augmentations are performed in a specific order (as shown below). Not all the steps will be certainly conducted, with each step having a probability of being chosen or dropped.

1. Random mask. We randomly replace some gene expression values with zero in the profile of the target cell. The mask percentage is 0.2, while the probability of executing the step is 0.5. Notice that this synthetic noise is similar to the dropout events in the single-cell sequencing experiments.
2. Gaussian noise. We randomly replace some gene expression values in the target cell profile with numbers drawn from a predefined Gaussian distribution. The noise percentage is 0.8. The mean of Gaussian distribution is 0, while the standard deviation is 0.2. The probability of executing this step is 0.5.
3. Random swap. For a gene expression profile, we randomly choose an even number of gene expression values and construct pairs from the subset, then swapping the gene expression values inside each pair. The total percentage that performs swapping is 0.1. The probability of execution is 0.5.
4. Crossover with another cell. We randomly choose another cell in the dataset as the crossover source and then select some genes from the target gene expression profile, swapping the gene expression value between the two cells. 25% of the gene expression data in one cell will be exchanged with the other cell. The probability of executing this step is 0.5. This exchanging step will not influence the next batch or the next training epoch.
5. Crossover with many cells. We randomly choose several cells in the dataset as the crossover source and some genes from the target gene expression profile, swap the expression values between the source cell and the target cells. 25% of the gene expression data in the cell will be exchanged with the selected cells. The probability of execution is 0.5. This step would not influence the next batch or the next training epoch.

### Architecture and hyperparameters

The encoder neural network in CLEAR consists of two fully connected layers. The query encoder and the key encoder share the same architecture. The first layer has 1024 nodes, while the second layer has 128 nodes. The *ReLU* function, defined as *ReLU*(*x*) = *max*(0, *x*), is used as the nonlinear activation function after the linear transformation. We use Adam optimizer with the learning rate as 1 and the cosine learning schedule. We train the paired neural networks for 200 epochs. Temperature, *τ*, in the CLEAR’s objective function, is set to be 0.2. The momentum coefficient *m* is 0.999. The hyper-parameters are determined using grid search with cross-validation.

### Performance evaluation

To evaluate the standard clustering performance of the proposed method, we use the adjusted Rand index (ARI) and Normalized Mutual Information (NMI). On the other hand, to benchmark different methods’ performance on batch effect removal, we utilize ARI and Average Silhouette Width (ASW). In addition, we also used cell ARI (cARI) and batch ARI (bARI), as well as cell ASW (cASW) and batch ASW (bASW). Their definitions are as follows. Note that, during evaluation, we use the default parameters for all the criteria. More quantitative measurement also shown in **Supplementary Method 3**.

ARI measures the similarity between two partitions by comparing all the pairs of the samples adjusted by random permutation.

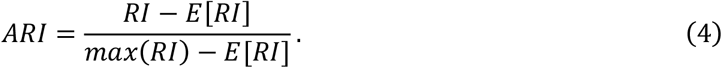

Where Rand index (RI) is defined as

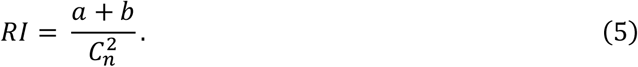

Where *a* is the number of pairs that are correctly labelled in the same set of the two partitions, and *b* is the number of pairs that are correctly labelled but not in the same set of the two partitions. 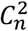 is the total number of pairs. *E*[*RI*] is the expected *RI* from a random model. ARI ranges from -1 to 1. A value close to 0 suggests random labeling, while close to 1 means the nearly perfect cell type purity. To evaluate batch effect removal, we also calculate three specific kinds of ARI, cell ARI (cARI) and batch ARI (bARI), and the Harmonic mean of the two f1_ARI. A higher cARI corresponds to higher cell type purity, while a bARI close to zero suggests superior batch effects removal.

NMI measures the amount of information obtained about one partition through observing the other partition, ignoring the permutations:

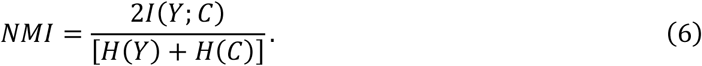

Where *Y* is the class labels, and *C* is the cluster label. *H*(.) is the entropy, and *I*(*Y*; *C*) measures the mutual information between *Y* and *C*.

ASW measures the relative distance between inter-clusters and intra-clusters. The Silhouette width (SW) is defined as:

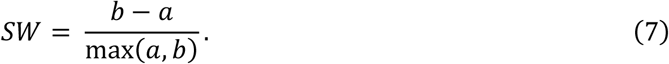

Where *a* is the mean distance between a sample and all the inter-cluster points, while *b* is the mean distance between a sample and all the other points in the next nearest cluster. The ASW is defined as the average of all the cells’ Silhouette width within the entire dataset. The range of ASW is [−1,1], where 1 suggests the best clustering result and -1 suggests the worst clustering result. To evaluate batch effect removal, we calculate three kinds of ASW, cell ASW (cASW), batch ASW (bASW), and the Harmonic mean of the two f1_ASW. A higher cASW suggests better cell type purity, while a lower bASW suggests better batch mixing.

### Software comparison and settings

To evaluate the performance of CLEAR compared with other methods, we select the below several software packages for comparison. All the evaluation codes and input data follow the instruction and tutorials provided by each package (**Code Availability**).

For baseline clustering, we compare CLEAR with R-based tools including Seurat, SC3, CIDR, SINCERA, scDHA, and SIMLR, and Python-based packages, such as ItClust, scVI, and scGNN. The details of the software we used are: (i) Seurat version 4.0.1 from CRAN. The parameters are set as the default value provided by the tutorial. (ii) SC3 version 1.2 from Bioconductor. The key parameter, svm_num_cells, which means the number of randomly selected training cells to be used for SVM prediction, is set as 5000. All other parameters follow the SC3 function instruction. (iii) CIDR version 0.1.5 from Github (https://github.com/VCCRI/CIDR). We set all the parameters as default following the README file on Github. (iv) SINCERA from Github (https://github.com/xu-lab/SINCERA). The parameters follow the pipeline of the demonstrations listed on their Github. (v) scDHA from Github (https://github.com/duct317/scDHA). Parameters and data input format are set following the running example. We also utilize its built-in function to generate 2d visualization of the representations. (vi) ItClust from Github (https://github.com/jianhuupenn/ItClust), running with the default parameters. For the sake of simplicity and convenience, we set the required source dataset in the code as “baron-human” across all the experiments. (vii) SIMLR version 1.18.0 from Bioconductor, running with the default parameters. We select the small-scale version of code because the large-scale version cannot run in our environment. (viii) scVI from Github (https://github.com/YosefLab/scvi-tools). We use K-means for clustering based on the embeddings generated from the trained VAE. (ix) scGNN version 1.0.2 from Github (https://github.com/juexinwang/scGNN). We run the GPU version and set the hyper-parameters following their example. We include LTMG inferring in preprocessing with the corresponding given option of the code. All the hyper-parameters are set following the tutorial.

For the dropout clustering experiment, we create a dropout_sampling function for random sampling. Random seeds are set to ensure each sampling is unique. Software parameters are the same as the baseline clustering experiment.

Seurat, SC3, CIDR, and SINCERA are run on the PC with Intel(R) Core i7-8750H CPU, Window 10 operation system, 32GB physical memory. The virtual memory limitation of our working environment is set as 100GB RAM, R version 4.0.3. We run scDHA, SIMLR, ItClust, scVI, and scGNN on a workstation with Intel(R) Xeon(R) CPU E5-2667 v4, CentOS Linux release 7.7.1908 operation system, Nvidia TITAN X GPU, 503GB physical memory.

### Case study of the COVID dataset

We first apply CLEAR on the published COVID PBMC scRNA-seq dataset, the parameters of CLEAR are set as in the **Architecture and hyperparameters**. Based on the 128 features generated by CLEAR, we run Seurat (V 4.0.1) with the parameter *Resolution* = 1.2 to cluster all the cells, by which 32 clusters are identified. The cell type of each cluster is annotated by the top differentially expressed genes found by the FindAllMarkers function. The statistical method is Wilcoxon Rank-Sum Test, and the *LogFC threshold* = 0.25. All the gene expression level plots are generated by the FeaturePlot function with the default parameters. To conduct different expression gene (DEG) analysis, we use the Wilcoxon Rank-Sum Test to search for the DEGs between each pair of monocytes obtained from the three groups (i.e., the health donors (HDs), moderate and severe (ARDS) groups). We put *LogFC threshold* = 0.25 and show negative (downregulated) genes as well. We obtain two groups of DEGs for each monocyte subtype and show the result in the Supplementary material (**Supplementary Table 4-5**). Given that gene expression score can be calculated by the AddModuleScore function, we use the function to calculate ISG score and MHC score for monocytes with the pre-determined interferon-stimulated gene set and MHC-related gene set (**Supplementary Table 6**). The significant test is also the Wilcoxon test in **Fig 5. g** and **h**. Finally, we use the DEGs we got before to perform Gene Ontology (GO) analysis for each COVID stage and run GSEA analysis on the GO results. All the parameters are set as default during GO and GSEA analysis.

## Data Availability

We used 10 datasets for evaluating the performance of clustering and dropouts, one dataset for benchmarking the batch effects removal. Two COVID-PBMC dataset for case study. The details information and the links to the publicly available sources of the 13 datasets can be found in the Method part.

## Code Availability

An open-source implementation of CLEAR is available at GitHub: https://github.com/ml4bio/CLEAR, under the MIT license.

## Contributions

### Competing Interests

The authors declare no competing interests.

**Additional Information**

## Notes

### Competing Interest Statement

The authors have declared no competing interest.

## References

1. Shapiro, E., Biezuner, T. & Linnarsson, S. Single-cell sequencing-based technologies will revolutionize whole-organism science. Nat Rev Genet 14, 618–630 (2013).

2. Shalek, A.K. et al. Single-cell RNA-seq reveals dynamic paracrine control of cellular variation. Nature 510, 363–369 (2014).

3. Maynard, A. et al. Therapy-Induced Evolution of Human Lung Cancer Revealed by Single-Cell RNA Sequencing. Cell 182, 1232–1251 e1222 (2020).

4. van Galen, P. et al. Single-Cell RNA-Seq Reveals AML Hierarchies Relevant to Disease Progression and Immunity. Cell 176, 1265–1281 e1224 (2019).

5. Tian, T., Zhang, J., Lin, X., Wei, Z. & Hakonarson, H. Model-based deep embedding for constrained clustering analysis of single cell RNA-seq data. Nat Commun 12, 1873 (2021).

6. Stegle, O., Teichmann, S.A. & Marioni, J.C. Computational and analytical challenges in single-cell transcriptomics. Nat Rev Genet 16, 133–145 (2015).

7. Kiselev, V.Y., Andrews, T.S. & Hemberg, M. Challenges in unsupervised clustering of single-cell RNA-seq data. Nature Reviews Genetics 20, 273–282 (2019).

8. Lopez, R., Regier, J., Cole, M.B., Jordan, M.I. & Yosef, N. Deep generative modeling for single-cell transcriptomics. Nature Methods 15, 1053-+ (2018).

9. Deng, Y., Bao, F., Dai, Q.H., Wu, L.F. & Altschuler, S.J. Scalable analysis of cell-type composition from single-cell transcriptomics using deep recurrent learning. Nature Methods 16, 311-+ (2019).

10. Satija, R., Farrell, J.A., Gennert, D., Schier, A.F. & Regev, A. Spatial reconstruction of single-cell gene expression data. Nat Biotechnol 33, 495–502 (2015).

11. Kiselev, V.Y. et al. SC3: consensus clustering of single-cell RNA-seq data. Nat Methods 14, 483–486 (2017).

12. Lin, P.J., Troup, M. & Ho, J.W.K. CIDR: Ultrafast and accurate clustering through imputation for single-cell RNA-seq data. Genome Biol 18, 59 (2017).

13. Guo, M., Wang, H., Potter, S.S., Whitsett, J.A. & Xu, Y. SINCERA: A Pipeline for Single-Cell RNA-Seq Profiling Analysis. PLoS Comput Biol 11, e1004575 (2015).

14. Wolf, F.A., Angerer, P. & Theis, F.J. SCANPY: large-scale single-cell gene expression data analysis. Genome Biol 19, 15 (2018).

15. Levine, J.H. et al. Data-Driven Phenotypic Dissection of AML Reveals Progenitor-like Cells that Correlate with Prognosis. Cell 162, 184–197 (2015).

16. Wang, B., Zhu, J., Pierson, E., Ramazzotti, D. & Batzoglou, S. Visualization and analysis of single-cell RNA-seq data by kernel-based similarity learning. Nat Methods 14, 414–416 (2017).

17. Park, S. & Zhao, H.Y. Spectral clustering based on learning similarity matrix. Bioinformatics 34, 2069–2076 (2018).

18. Butler, A., Hoffman, P., Smibert, P., Papalexi, E. & Satija, R. Integrating single-cell transcriptomic data across different conditions, technologies, and species. Nat Biotechnol 36, 411–420 (2018).

19. Dijk, D.v. et al. Recovering Gene Interactions from Single-Cell Data Using Data Diffusion. Cell 174, 716-729.e727 (2018).

20. Eraslan, G., Simon, L.M., Mircea, M., Mueller, N.S. & Theis, F.J. Single-cell RNA-seq denoising using a deep count autoencoder. Nature Communications 10, 390 (2019).

21. Tran, D. et al. Fast and precise single-cell data analysis using a hierarchical autoencoder. Nature Communications 12, 1029 (2021).

22. Hu, J. et al. Iterative transfer learning with neural network for clustering and cell type classification in single-cell RNA-seq analysis. Nat Mach Intell 2, 607–618 (2020).

23. Wang, J. et al. Data denoising with transfer learning in single-cell transcriptomics. Nat Methods 16, 875–878 (2019).

24. Wang, J.X. et al. scGNN is a novel graph neural network framework for single-cell RNA-Seq analyses. Nature Communications 12, 1882 (2021).

25. Li, X.J. et al. Deep learning enables accurate clustering with batch effect removal in single-cell RNA-seq analysis. Nature Communications 11 (2020).

26. Ding, J.R., Condon, A. & Shah, S.P. Interpretable dimensionality reduction of single cell transcriptome data with deep generative models. Nature Communications 9 (2018).

27. Ding, J. & Regev, A. Deep generative model embedding of single-cell RNA-Seq profiles on hyperspheres and hyperbolic spaces. Nat Commun 12, 2554 (2021).

28. Pierson, E. & Yau, C. ZIFA: Dimensionality reduction for zero-inflated single-cell gene expression analysis. Genome Biol 16 (2015).

29. Risso, D., Perraudeau, F., Gribkova, S., Dudoit, S. & Vert, J.P. A general and flexible method for signal extraction from single-cell RNA-seq data. Nat Commun 9, 284 (2018).

30. Brbic, M. et al. MARS: discovering novel cell types across heterogeneous single-cell experiments. Nat Methods 17, 1200–1206 (2020).

31. Chen, T., Kornblith, S.M.N., & Hinton, G. A Simple Framework for Contrastive Learning of Visual Representations. ICML-2020 (2020).

32. Deng, Q., Ramsköld, D., Reinius, B. & Sandberg, R. Single-cell RNA-seq reveals dynamic, random monoallelic gene expression in mammalian cells. Science 343, 193–196 (2014).

33. Yan, L. et al. Single-cell RNA-Seq profiling of human preimplantation embryos and embryonic stem cells. Nature structural & molecular biology 20, 1131–1139 (2013).

34. Pollen, A.A. et al. Low-coverage single-cell mRNA sequencing reveals cellular heterogeneity and activated signaling pathways in developing cerebral cortex. Nature biotechnology 32, 1053–1058 (2014).

35. Kolodziejczyk, A.A. et al. Single cell RNA-sequencing of pluripotent states unlocks modular transcriptional variation. Cell stem cell 17, 471–485 (2015).

36. Muraro, M.J. et al. A single-cell transcriptome atlas of the human pancreas. Cell systems 3, 385-394. e383 (2016).

37. Hrvatin, S. et al. Single-cell analysis of experience-dependent transcriptomic states in the mouse visual cortex. Nature neuroscience 21, 120–129 (2018).

38. Consortium, T.M. A single cell transcriptomic atlas characterizes aging tissues in the mouse. Nature 583, 590 (2020).

39. Luecken, M.D. et al. Benchmarking atlas-level data integration in single-cell genomics. BioRxiv (2020).

40. Tran, H.T.N. et al. A benchmark of batch-effect correction methods for single-cell RNA sequencing data. Genome biology 21, 1–32 (2020).

41. Wolf, F.A. et al. PAGA: graph abstraction reconciles clustering with trajectory inference through a topology preserving map of single cells. Genome biology 20, 1–9 (2019).

42. Trapnell, C. et al. The dynamics and regulators of cell fate decisions are revealed by pseudotemporal ordering of single cells. Nature biotechnology 32, 381–386 (2014).

43. Wilk, A.J. et al. A single-cell atlas of the peripheral immune response in patients with severe COVID-19. Nat Med 26, 1070–1076 (2020).

44. Kuri-Cervantes, L. et al. Immunologic perturbations in severe COVID-19/SARS-CoV-2 infection. bioRxiv (2020).

45. Kuri-Cervantes, L. et al. Comprehensive mapping of immune perturbations associated with severe COVID-19. Sci Immunol 5 (2020).

46. Zhao, J. et al. Antibody Responses to SARS-CoV-2 in Patients With Novel Coronavirus Disease 2019. Clinical Infectious Diseases 71, 2027–2034 (2020).

47. Choudhary, S., Sharma, K. & Silakari, O. The interplay between inflammatory pathways and COVID-19: A critical review on pathogenesis and therapeutic options. Microb Pathog 150, 104673 (2021).

48. Hu, B., Huang, S. & Yin, L. The cytokine storm and COVID-19. Journal of Medical Virology 93, 250–256 (2021).

49. Schulte-Schrepping, J. et al. Suppressive myeloid cells are a hallmark of severe COVID-19. medRxiv, 2020.2006.2003.20119818 (2020).

50. Unterman, A. et al. Single-Cell Omics Reveals Dyssynchrony of the Innate and Adaptive Immune System in Progressive COVID-19. medRxiv, 2020.2007.2016.20153437 (2020).

51. Guo, C. et al. Single-cell analysis of two severe COVID-19 patients reveals a monocyte-associated and tocilizumab-responding cytokine storm. Nat Commun 11, 3924 (2020).

52. Ragab, D., Salah Eldin, H., Taeimah, M., Khattab, R. & Salem, R. The COVID-19 Cytokine Storm; What We Know So Far. Frontiers in Immunology 11 (2020).

53. Schulte-Schrepping, J. et al. Severe COVID-19 Is Marked by a Dysregulated Myeloid Cell Compartment. Cell 182, 1419–1440 e1423 (2020).

54. Deng, Y., Bao, F., Dai, Q., Wu, L.F. & Altschuler, S.J. Scalable analysis of cell-type composition from single-cell transcriptomics using deep recurrent learning. Nat Methods 16, 311–314 (2019).

55. Villani, A.-C. et al. Single-cell RNA-seq reveals new types of human blood dendritic cells, monocytes, and progenitors. Science 356 (2017).

56. Ren, X. et al. COVID-19 immune features revealed by a large-scale single-cell transcriptome atlas. Cell 184, 1895-1913. e1819 (2021).

57. Chen, X., Fan, H., Girshick, R. & He, K. Improved baselines with momentum contrastive learning. arXiv preprint 2003.04297 (2020).

